# Probing glioblastoma and its microenvironment using single-nucleus and single-cell sequencing

**DOI:** 10.1101/775197

**Authors:** Sen Peng, Sanhita Rath, Connor Vuong, Saumya Bollam, Jenny Eschbacher, Xishuang Dong, Shwetal Mehta, Nader Sanai, Michael Berens, Seungchan Kim, Harshil Dhruv

## Abstract

Single-cell (scSeq) and single-nucleus sequencing (snSeq) are powerful tools to investigate cancer genomics at single cell resolution. Multiple studies have recently illuminated intratumoral heterogeneity in glioblastoma, however, the majority focused on molecular complexity of tumor cells, without considering unexplored host cell types that contribute to the microenvironment around tumor. To address the glioblastoma microenvironment composition and potential tumor-host interactions, we performed deep coverage sequencing of freshly resected primary GBM patient tissue without implementing any tumor enrichment strategies. The sequencing resulted in 902 cells and 1186 nuclei, respectively, passing quality control and with low mitochondrial gene percentage. We customized reference transcriptome by listing gene transcript loci as exons to take into account immature RNA, which greatly improved the alignment rate for single-nucleus data. We applied Cell Ranger pipelines (Version 3.0.2) and Seurat package (Version 2.3.1) and discovered 10 clusters in both scSeq and snSeq. Pathway analysis of each cluster signature in scSeq data along with known GBM microenvironment cell signatures revealed glioma tumor population along with surrounding microglia/macrophages, astrocytes, pericytes, oligodendrocytes, T cells and endothelial cells. The analysis of snSeq was able to capture the majority of cell types from patient tissues (tumor and microenvironment cells), but interestingly presented different cell type composition in microenvironment cell types such as microglia/macrophages. Integrating single-cell and single-nucleus transcriptomic data using canonical correlation analysis facilitated a comparison of snSeq and scSeq, contrasting depiction for certain cell types (e.g. NKX6-2 gene in Oligodendrocytes). Differential analysis of pathways between tumor and microenvironment cells unveiled potentially rewired pathways such as double strand break repair pathway. Our results demonstrate the cellular diversity of brain tumor microenvironment and lay a foundation to further investigate the individual tumor and host cell transcriptomes that are influenced not only by their cell identity but also by their interaction with surrounding microenvironment.

## I. Introduction

Single-cell (scSeq) and single-nucleus sequencing (snSeq) has shed light on cancer genomics at single cell resolution including the central nervous system (CNS) [1, 2]. Although bulk tumor sequencing can extend our understanding of the cellular states and functions of tumor cells, bulk measurements mix the signal of the diverse cells within each tumor and sometimes along with its microenvironment cells, thus masking potential vital differences and only providing convoluted insight into tumor cell development and tumor microenvironment (TME) influences. Single-cell (scSeq) and single-nucleus sequencing (snSeq) can help tackle these limitations of bulk sequencing and are powerful tools to investigate brain tumor and its microenvironment genomics at unprecedented single cell resolution.

Cancer could be regarded as a complex, heterogeneous and evolutionary process. Intra-tumor heterogeneity (ITH), both at genetic and transcriptomic levels, is one of the hallmarks of brain cancer and contributes to therapy resistance [3]. Single-cell RNA-seq provides the opportunity to generate the detailed transcriptomic cell profiling required to address such tumor heterogeneity. Underlying molecular biology and chemistry of single-cell library preparation have improved considerably from the initial proof-of-principle studies [4, 5]. Accompanying the enormous progress of multiple molecular layers of single-cell sequencing technologies, computational methods and bioinformatic algorithms have also been developed to best process and interpret the single-cell data for normalization, feature selection, differential gene expression analysis, clustering and trajectory analysis [6, 7]

Despite the tremendous advances, certain limitations and challenges still exist in single-cell sequencing. First, current single cell approaches still require live cells and are incompatible with frozen archival samples, which complicates studies when tissue availability is unpredictable or the studies involve multi-institute collaboration. Second, single cell protocols might be biased toward particular cell types, since dissociation method may destroy sensitive cells while failing to separate cells that are surrounded by brain extracellular matrix [8]. Finally, the enzymatic and mechanical dissociation processes used to isolate single cells has been shown to influence the transcriptomic profile of the cells and could introduce stress-related transcriptional artifacts [9].

Hence, single nuclei isolation has evolved as an alternative method for various transcriptomic studies, especially when studying highly interconnected tissues, such as glioblastoma multiforme (GBM). Single nucleus sequencing is free of both RNA degradation and artificial transcriptional stress responses, as it circumvents enzymatic dissociation for nuclear isolation, and therefore maintains transcriptome integrity. Studies of mouse visual cortex cells demonstrated that only small differences lie between total cellular and nuclear transcriptome [9]. snSeq could provide wider sample applicability (fresh and archival), reduced dissociation bias and comparable gene detection compared to scSeq.

Besides the genetic, transcriptomic and epigenetic heterogeneity among tumor clones, heterogeneity among tumor-infiltrating cells in the microenvironment also plays essential roles in tumor evolution, invasion, immune response, metastasis, and resistances to various therapies [10]. In addition to tumor cells themselves, brain tumors are also infiltrated with other cells such as endothelial cells, pericytes, fibroblasts, and immune cells. Thorough understanding of the composition, interactions, and dynamics of cancer ecosystems is key to understanding tumor fitness, evolution, and the emergence of therapy resistance. Computational deconvolution approaches could help infer the mixed cellular composition of brain tumors along with its microenvironment, but such analyses are limited to a few known cell types and sensitive to parameter estimation due to its inference nature [11, 12]. Moreover, in silico deconvolution often requires characterization of pure known cell types and is not always available. Single-cell and nucleus sequencing represent a monumental technological leap, since it allows precise dissection of the complex ecosystems of tumors while capturing rare cell types.

In this study, we performed deep coverage bulk, single cell, and single nucleus sequencing of freshly resected primary GBM patient tissue without implementing any tumor enrichment strategies. It enables us to investigate the feasibility of snSeq, compare the single cell with single nucleus data and probe into the glioblastoma microenvironment composition and potential tumor-host interactions.

## II. Methods

### A. Single cell isolation and sequencing

Single cell RNA-Seq libraries from the fresh sample was prepared using GemCode single Cell 3’ Gel Bead and Chromium™ Single Cell 3’ Library Kit (10x Genomics, CA) as per manufacturers protocol. The library was then sequenced with Illumina NextSeq system. We aimed to sequence ~2,000 cells per sample and to achieve sequencing depth of 100,000 reads per cell. Quantification of cDNA libraries was performed using Qubit dsDNA HS Assay Kit (Life Technologies).

### B. Single nucleus isolation and sequencing

Brain tissue samples were partially minced and frozen in DMSO solution before nuclei extraction and library preparation. The brain tissue was centrifuged at 300 g for 2-3 mins and colored DMSO was carefully discarded. The pellet was washed with 1 mL of PBS + 1% BSA pH 7.4 and centrifuged at 300 g for 2-3 min. The pellet was resuspended in 1.5 mL of HB buffer (Combine 1.5 mL of NIM2 buffer with 150 uL of 1% NP40, 15 uL of SuperaseIN RNAase inhibitor, 15 uL of RNaseIN inhibitor, and 10 uL DNase I stock solution) and then transferred to glass homogenization tube using cut tip. The sample was minced with 5-7 strokes of Loose Pestle (A) and 10-15 strokes of Tight Pestle (B). The sample was then incubated on ice for 10 mins before filtered through a 40-um cell strainer. After Centrifuging for 8 min at 300 g, the pellet was resuspended in 1 mL of chilled PBS + 1% BSA pH 7.4 and pipetting up and down 10 times was conducted to homogenize the sample. The sample was filtered through a 20-um cell strainer and then centrifuged for 8 mins at 300 g. 20 uL of NucGreen dye along with 600 uL of PBS + 1% BSA pH 7.4 buffer were added to the sample, and shake to mix, and briefly spin down.

FACS sorting was performed using SH800 Flow Cytometry Apparatus with the following steps: 1) Set sample pressure to 6 and adjust temperature in sample and flow cell chambers to 4°C prior to start. 2) First record populations for non-stained sample and compare to stained sample to determine the location of the nuclei population under FITC vs SSC parameters. 3) Sort nuclei into 150 uL PBS + 1% BSA or into 3’ and 5’ library preparation master mixes until sample depletion (or until desired nuclei count for sequencing). RNA was subsequently extracted with the help of DNase I in RNase-free water. Quality check resulting RNA using High Sensitivity RNA TapeStation and followed immediately by the 10x Genomics Single Cell Protocol to generate sequencing library.

### C. Single cell/nucleus analysis and statistics

Raw sequencing data were preprocessed using CellRanger 3.0.2. In brief, Cellranger mkfastq module was applied to generate fastq files by demultiplexing Chromium-prepared sequencing samples based on their barcodes. Those fastq files were then input to cellranger count to generate UMI count data at a single-cell resolution. Further single-cell data analysis was conducted using R package Seurat. In brief, we applied initial normalization where UMI counts for each cell were scaled by total expression and a factor equal to the median counts of all genes. Data regression was performed using the ScaleData function with nUMI, mitochondrial read percentage as confounding factors.

Cells expressing more than 6500 genes were filtered out for potential cell aggregates. Cells with a percentage of mitochondrial genes expression > 0.1 were also filtered out for probable dead cells. These expression values were log transformed before further downstream analyses. Principle component analysis, variable gene identification, Shared Nearest Neighbor (SNN) clustering analysis, and t-distributed stochastic nearest neighbor embedding (tSNE) visualization were then performed. In details, the first 10 principle components were used for clustering analysis and clusters were visualized with tSNE mapping. Signature markers for each specific cluster were identified with function FindAllMarkers against all remaining cells. Top 100 significant markers with largest average log fold change were retained as signature for each cluster.

We determined the brain cell types in each of the cluster by evaluating those makers along with the expression of known signatures genes for brain cell types including astrocytes, oligodendrocytes, microglia, pericytes, macrophages, T cells and endothelial cells [13, 14]. Pseudotime trajectory analysis was conducted using R package monocle (version 2.8.0) with author recommended default settings [6]. Genes used for trajectory ordering were taken from the dispersion genes with normalization log2 mean expression > 0.1. DDRTree method was used for dimension reduction and cell ordering along the single-cell trajectories.

Gene set variation analysis (GSVA) [15] was performed to determine the activities of GBM molecular subtype [16] signatures in each cell’s transcriptome data. Cells with highest subtype signature GSVA score were classified to the corresponding subtype. All statistical tests and figures were generated using various packages in R 3.6.1.

### D. Evaluation of Differential Dependency (EDDY) analysis

Evaluation of Differential DependencY (EDDY) [18] is a statistical gene set test method to detect differential genetic dependencies between conditions in order to better understand underlying molecular features and their mechanisms. Specifically, EDDY evaluates the probability distributions of dependency gene networks, which is different from differential expression of individual genes or correlation changes of individual gene-gene interactions (See Figure 5a for overview of the algorithm). When compared to Gene Set Co-expression Analysis (GSCA), EDDY generates lower false positives, where GSCA identifies differentially co-expressed gene sets by analyzing pair-wise gene-gene interactions. The Java implementation of EDDY is freely available to noncommercial users at http://biocomputing.tgen.org/software/EDDY.

In this study, we employed EDDY-GPU^1^, that is the GPU version of EDDY, as the number of cells becomes too big for the Java version of EDDY. We explored intra-tumoral heterogeneity of the tumor on the scRNA-seq data, by first identifying subpopulation of tumor cells and other non-tumor cells from its surrounding, i.e. microenvironment and analyzing differential gene dependencies across those subpopulations. The program was run on high-performance GPU clusters (NVIDIA Tesla P100) at PVAMU’s Advanced Computing Lab.

## III. Results

### A. Study design

GBM patient tumor along with its microenvironment was collected and divided into three parts. Two samples were flash frozen and stored for further process (including bulk and single nucleus sequencing), the third one was immediately dissociated and used to generate single cell library using 10X Genomics platform. Fig. 1 shows the scheme of the study design. The three sequencing technologies on the same patient tumor facilitate the comparisons among those approaches as well as the comprehensive understanding of the tumor biology.

**Fig. 1.**
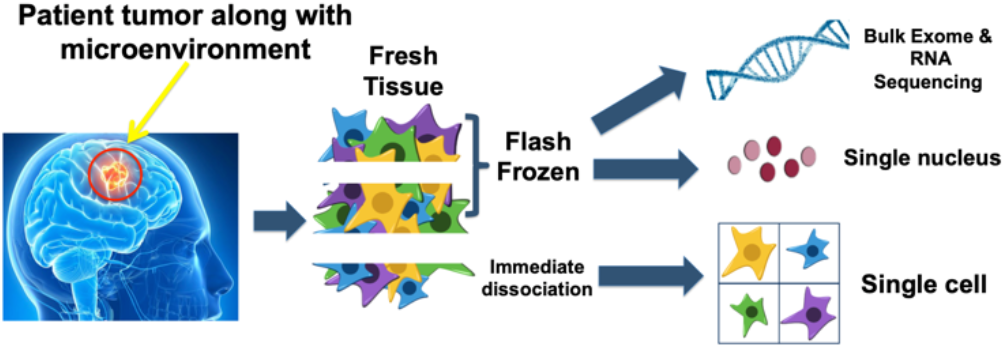
Scheme of study design

### B. Mapping the Transcriptional Landscape of glioblastoma patient tumor using single cell sequencing

To address the glioblastoma microenvironment composition and potential tumor-host interactions, we generated scSeq libraries of freshly resected primary GBM patient tissue without implementing any tumor cell enrichment strategies. Single cell sequencing libraries were prepared using 10X Chromium Gemcode machine and sequenced on Illumina NextSeq 500. Preliminary data analysis showed transcriptomic profile of 902 single cells at the deep coverage of 176,000 reads per cell from frozen GBM patient tissue. This run was of high quality with 2,663 median genes per cell and low mitochondrial gene percentage (median < 5%).

Single cell sequencing data were analyzed using the Cell Ranger analysis pipelines and Seurat packages. We identified 10 clusters with a recommended resolution parameter 0.6 and employed the TSNEPlot function to generate a visual representation of the clusters using T-distributed Stochastic Neighbor Embedding (tSNE) (Fig. 2a). Pathway analysis of each cluster signature along with known GBM microenvironment cell signatures revealed annotated brain cell types: Tumor cells (Ki67 +ve and Ki67-ve), Astrocytes, Oligodendrocytes, Antigen Presenting Cells (Macrophages and Microglia), Endothelial cells, Pericytes, T cells, and Erythrocytes (Fig. 2c). Expression of key signature genes related to specific cell types were visualized in Fig. 2b. Top three most differentially expressed genes in each identified cluster as compare to all other clusters are also shown here in the form of heatmap Fig. 3b (top). With the help of pseudotime analysis, the GBM brain cells was ordered along a trajectory, and cells at different states at one branching points were identified. Cluster 0&3 (tumor cells) were found at one end of the trajectory, and then followed the path of other microenvironment cell to the other end of cluster 1 (microglia/macrophages) (data not shown).

**Fig. 2.**
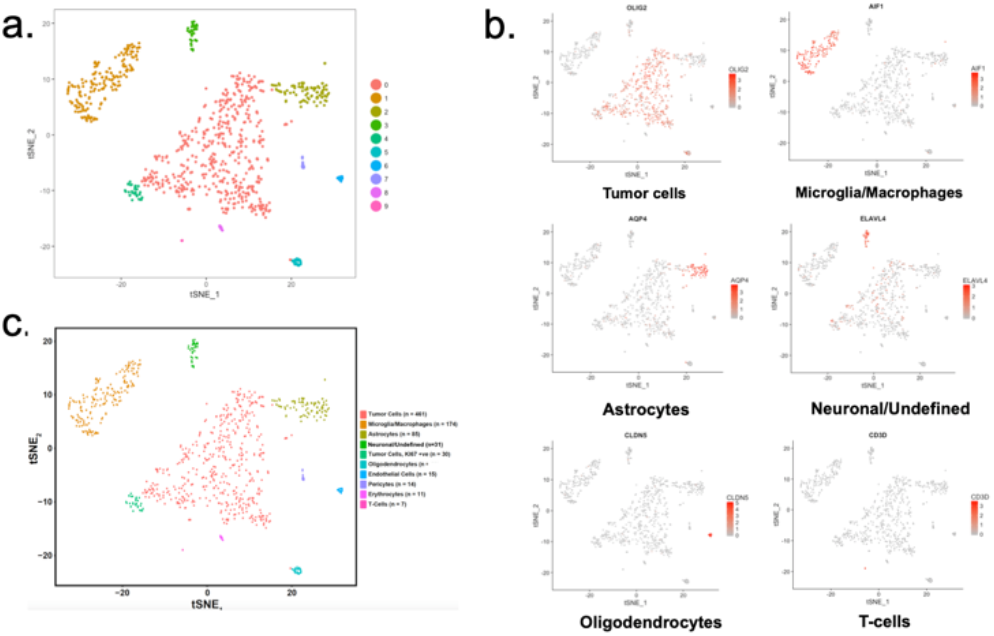
a. tSNE map of patient cells by clusters in scSeq: each dot represents individual cell and colors correspond to clusters. b. Oligo2 is highly expressed and localized in cluster 0, indicating tumor cell type. Other known signatures: AQP4 (astrocytes; C2), AIF1 (Microglia/Macrophages; C1), ELAVL4 (Neuronal/Undefined; C3), CLDN5 (Oligodendrocytes; C5) and CD3D (T cells; C9). c. Clusters were annotated by their signatures and known cell type markers.

**Fig. 3.**
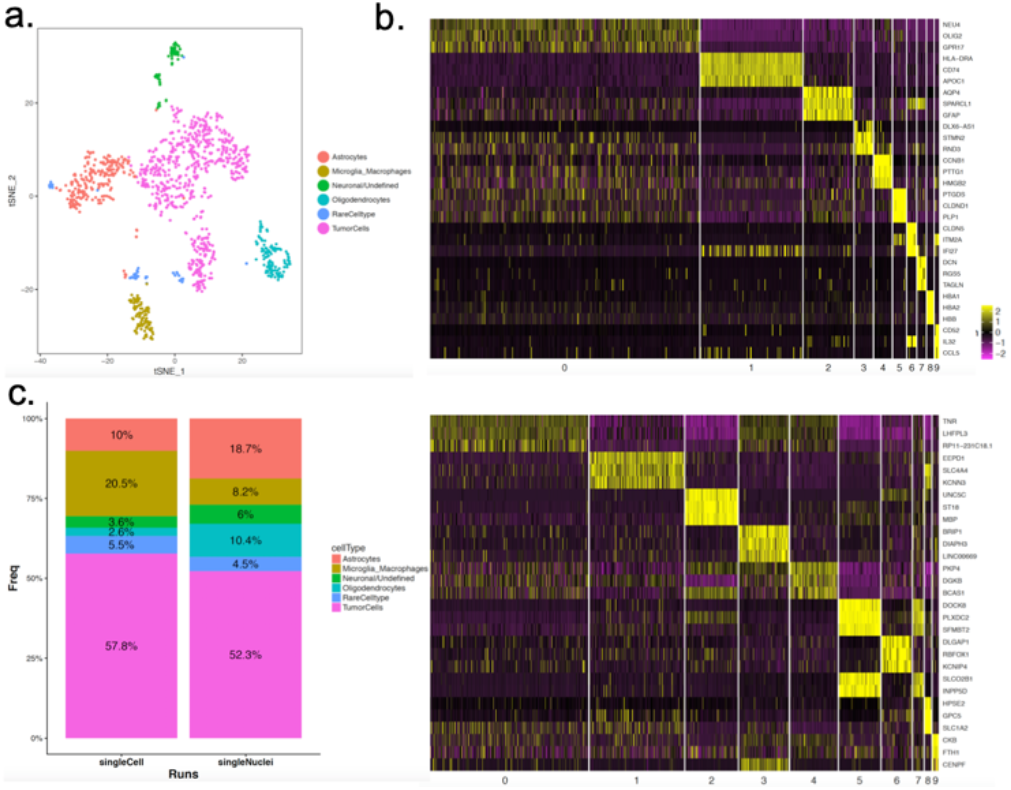
a. tSNE map with annotated clusters in snSeq. b. Cluster signatures: Heatmap showing top three most differentially expressed genes (DEG) across each cluster as compared to all other clusters (top: scSeq; bottom: snSeq). c. cell type distribution in a patient tumor.

### C. Single nucleus analysis and comparison to single cell data

To characterize the differences between nuclei and whole cells and the ability to detect cell types, snSeq library was prepared using aforementioned method targeting 2,000 cells and deep coverage sequencing was performed on Illumina NextSeq 500. Nuclei extraction protocol for human brain tissue were tested and modified to achieve good quality library and final protocol was described in method section. The CellRanger software was utilized to align the reads based on STAR aligner and quantify gene expression. By default, CellRanger quantifies expression for mature messenger RNA (mRNA) by counting reads aligned to exons as annotated in the human genome reference. However, the snSeq profiles nuclear precursor mRNA (pre-mRNA), which include transcripts that have not finished RNA splicing to be transformed into a mature messenger RNA. Intronic reads may also reflect cell type specific features, such as retained introns or alternative isoforms. To capture all the information in the pre-mRNA, we aligned the reads to a custom “pre-mRNA” reference that includes the intron region information. In this way, the intronic reads from pre-mRNA are included in the UMI counts for each gene and barcode. We aligned and quantified gene expression using both mature and pre-mRNA references, and these reads were further cleaned, QCed and compared.

We observed that when we aligned the reads to pre-mRNA reference, the number of genes called per nucleus increase significantly. For example, when using pre-mRNA as a reference, we observed a 18.3% increase in the number of the median UMI counts per nuclei (3118 vs 3689) and an 86.9% increase for the median genes per nuclei (1976 to 3693). Also, the percentage of reads mapped confidently to transcriptome (pre-mRNA reference) increased from 18.3% to 55.2%. Those results demonstrated that intronic region annotation is required for accurate gene quantification and downstream cell type identification from snSeq.

Although gene dropouts were higher in nuclei than in cells (mean genes detected 2663 cell vs 1976 nuclei), We could identify similar cell types using nuclei data as compared to whole cells (Fig. 3a). Rare cell types (<5% of population, including Pericytes and T cells) were not detected in nuclei data. This could be due to single nucleus library preparation, amplification stage efficiency, or detection power of sample size [17]. Overall, tumor cells were the largest portion and comprised of more than 50% of whole brain tissue in both single cell and nucleus data (Fig. 3c). This result demonstrated that using single nuclei to study brain tumor was relatively unbiased and could identify the major cell types. Integrating single-cell and single-nucleus transcriptomic data using canonical correlation analysis facilitated a comparison of snSeq and scSeq, contrasting depiction for certain cell types. For example, NKX6-2 gene expression was only detected in Oligodendrocytes in single cell data while not in single nucleus data.

In general, we observed comparable cell types identified with nuclei and cells and matched cluster proportions were mostly consistent, except that microglia/macrophage cells were less-represented among nuclei data (Fig. 3c). This could be due to GBM heterogeneity or nuclear content varies among cell types and for certain specific cell type, though previous study advocated that nuclear profiling is likely to be less cell type biased than scSeq [9].

### D. Molecular subtypes of glioblastoma tumor cells

To investigate the GBM heterogeneity at single cell resolution, Gene Set Variation Analysis (GSVA) was applied to determine the glioblastoma molecular subtype gene signature [16] score and each single cell/nucleus was classified according to their highest signature score. As shown in Fig. 4, both single cell and nucleus data indicated that the tumor (center cluster) is mostly comprised of mesenchymal type, which is consistent with bulk RNAseq data (data not shown). Those heterogenous cells of minor classical and proneural subtype appear closer in dimensional space to other microenvironment cells, indicating more similar transcriptome to surrounding cells compared to other majority tumor cells. This result illustrated the tumor-host interactions and bidirectional influence of tumor microenvironment.

**Fig. 4.**
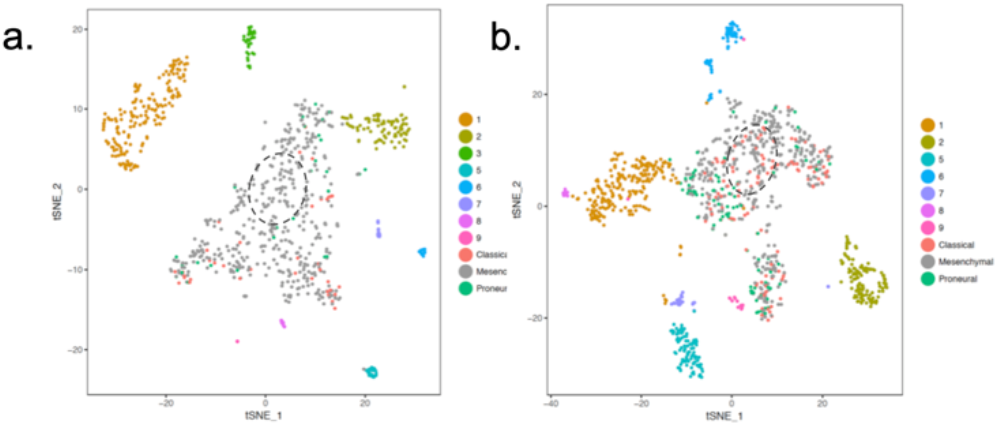
a. Single cell data: glioma tumor cells were classified to molecular subtypes. Dotted line indicates center region of majority mesenchymal subtype. b. Single nucleus data.

### E. Differential Dependency Analysis (EDDY) between tumor and microenvironment cells

Considering complex molecular mechanisms and heterogeneity of cancer, the discovery of biomarkers and subtype-specific drug targets must be based on network-driven activities of a gene set rather than individual genes. Such insight requires an understanding of gene interdependence rather than merely the more commonly utilized analyses of simple differential gene expression among comparative sets. We’ve applied EDDY-GPU as described previously to identify pathways enriched with differential dependencies in tumor and microenvironments, assisted by existing prior knowledge of gene interactions (Fig. 5a).

**Fig. 5.**
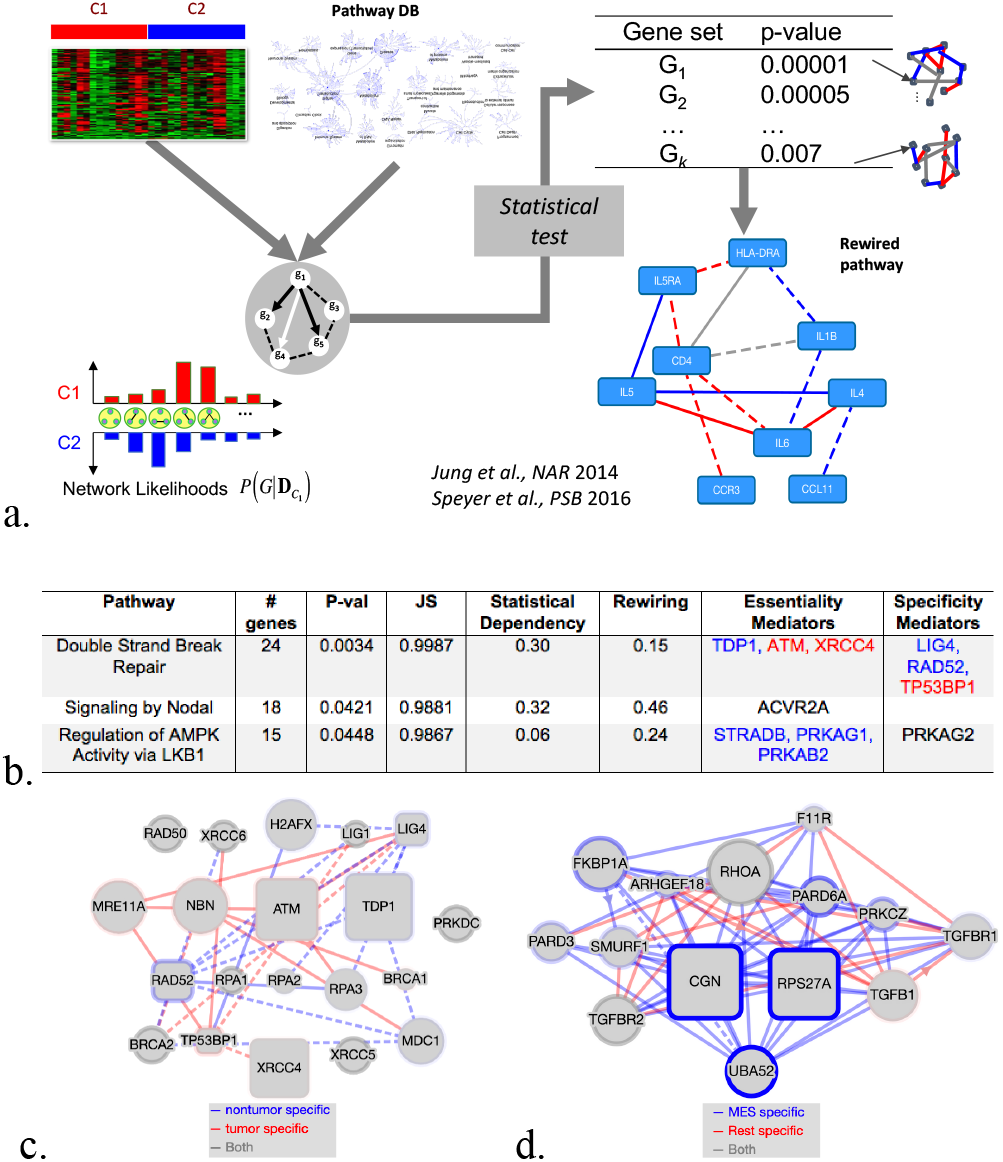
a. Interrogation of gene sets (pathways) for differential dependencies between tumor and microenvironment cells. b. Pathways rewired between GBM tumor cells and non-tumor cells.

Due to technological limit of current single cell RNA sequencing platform, the transcriptomic profile of each cell tends to be sparse, and our initial data showed 83.4% sparsity. In order to make the data more amenable for downstream analysis, including EDDY analysis, we averaged the detected transcript expression in each cell with its 3-nearest neighbors. This strategy greatly reduced the sparsity percentage from 83.4% to 64.9%. The merged data was then used to explore rewired pathways between different cell types and subtypes of GBM.

#### Tumor vs. Non-Tumor cells

We performed EDDY analysis of scSeq data from GBM patient, first by comparing tumor cells against non-tumor cells identified in the analysis described above, then, by comparing MES subtype of tumor cells and non-MES tumor cells. Before performing EDDY analyses, the raw counts of scSeq data were binarized, by converting non-zero counts to 1 and zero counts to 0. The threshold of zero was chosen considering the sparsity of raw counts in scSeq data.

Comparing the two groups of 419 tumor cells and 317 non-tumor cells, EDDY yielded 3 pathways enriched with differential dependency (Fig. 5b). The full results are also available at https://ccsb.pvamu.edu/eddy/NVIDIA/TvsNT/. Interestingly, EDDY analysis demonstrated differential dependency in DNA double strand break repair pathway (fig. 5c), which may play critical role in response to standard of care therapy, that is TMZ + Radiation. It will be interesting to study the roles of essentiality and specificity mediators identified by EDDY in mediating the tumor cell response and non-tumor cell response to standard of care therapy to identify potentially novel targets for improving the efficacy of standard of care therapy.

*GBM Mesenchymal vs. Non-Mesenchymal*: Using known molecular markers of GBM Mesenchymal subtype (50 genes) [16], we found most of our tumor cells be mesenchymal subtype (427 cells out of 491 tumor cells). Comparison between MES and non-MES tumor cells resulted in 14 pathways enriched with differential dependency. The full results are available online at https://ccsb.pvamu.edu/eddy/NVIDIA/MESvsRest. TGF beta has been shown to induce mesenchymal phenotype and tumor cell invasion in GBM [19]. EDDY analysis identified differential dependency in TGF beta receptor signaling (fig. 5d) in mesenchymal vs non-mesenchymal GBM cells. As part of future work, we would like to functionally perturb essentiality mediators of the TGF beta receptor signaling identified by EDDY and assess molecular subtype distribution of tumor cells. In addition to TGF beta receptor signaling pathway, lysosome vesicle biogenesis and gap junction degradation pathway also show differential dependency between mesenchymal and non-mesenchymal tumor cells, and may play important role in invasive behavior of mesenchymal glioma cells.

## IV. Conclusion

In conclusion, our findings indicate that single cell sequencing provides a valuable resource that can improve our understanding of the glioblastoma tumor along with its microenvironment. We also showed that single nucleus sequencing could be successfully applied to capture the majority cell types from GBM patient tissues (including both tumor and microenvironment cells), but with slightly different capture efficiency as compared to single cell sequencing. In general, deep snSeq is well suited for large-scale surveys of cellular diversity in brain tissue as it provides similar resolution for cell type detection to scSeq. Higher resolution of tumor subtype analysis revealed mixed subtype population, with heterogenous minor subtype cells tend to cluster closer to microenvironment cells compared to majority tumor cells, suggesting tumor host interactions. In addition, EDDY analysis revealed differential pathway dependencies between GBM tumor cells and microenvironment cells. Our analysis provides a general framework to decipher brain tumor cell genotypes and the composition of the TME. Our results demonstrate the cellular diversity of brain tumor microenvironment and lay a foundation to further investigate the individual tumor and host cell transcriptomes that are influenced not only by their cell identity but also by their interaction with surrounding microenvironment.

## Footnotes

1 https://github.com/dolchan/eddy-gpu

